# Non-histone protein methylation in *Trypanosoma cruzi* epimastigotes

**DOI:** 10.1101/2022.03.12.484072

**Authors:** Rafael Fogaça de Almeida, Aline Castro Rodrigues Lucena, Michel Batista, Fabricio Klerynton Marchini, Lyris Martins Franco de Godoy

**Author notes:** Corresponding author: Lyris Martins Franco de Godoy; Rua Prof. Algacyr Munhoz Mader, 3775, CIC 81350-010 Curitiba/PR, Brasil.

## Abstract

Post-translational methylation of proteins, which occurs in arginines and lysines, modulates several biological processes at different levels of cell signaling. Recently, methylation has been demonstrated in the regulation beyond histones, for example, in the dynamics of protein-protein and protein-nucleic acid interactions. However, the presence and role of non-histone methylation in *Trypanosoma cruzi*, the etiologic agent of Chagas disease, has not yet been elucidated. Here, we applied mass spectrometry-based-proteomics (LC-MS/MS) to describe the methylproteome of *T. cruzi* epimastigotes. A total of 1253 methyl sites in 824 methylated proteins were identified, of these, 712 (86.4%) proteins were identified in two or more biological replicates. Our results of functional enrichment analysis, protein-protein interaction, and co-occurrence with other PTMs, show that protein methylation impacts different levels of biological processes, such translation, RNA and DNA binding, amino acid and carbohydrate metabolism. In addition, 171 methylated proteins were previously reported with phosphorylation sites in *T. cruzi*, including flagellar proteins and RNA binding proteins, indicating that there may be an interplay between these different modifications in non-histone proteins. Our results show that a broad spectrum of functions are affected by methylation indicating its potential to impact important processes in the biology of the parasite and other trypanosomes.

**Statement of the significance:** *Trypanosoma cruzi* is a protozoan parasite that causes Chagas’ disease in humans and faces different environmental changes throughout its life cycle. Protein methylation is an important post-translational modification by which cells respond and adapt to the environment. We applied mass spectrometry-based proteomics and reported the first proteomic analysis of arginine and lysine methylation in *T. cruzi*. Our data demonstrate that protein methylation is broad and impacts different cellular processes in *T. cruzi*. This study represents a significant advance in the investigation of the importance of protein methylation in trypanosomes.

## 1. Introduction

Post-translational modifications (PTMs) impact different biological functions and significantly contribute to cellular homeostasis and environment adaptation^[1]^. Among them, an important PTM is protein methylation^[2,3]^ which occurs in lysine and arginine residues and impacts fundamental cellular events, from gene transcription and RNA processing to protein translation and cell signaling^[4]^.

Arginine and lysine methylation are catalyzed by S-adenosylmethionine (SAM)-dependent protein arginine methyltransferases (PRMTs) and protein lysine methyltransferases (PKMTs), respectively^[5]^. In humans, the most common PRMT is type I, which catalyzes the formation of monomethyl-arginine (MMA/Rme1), transferring a methyl group from SAM to guanidine nitrogen. It can add a second methyl group at the same nitrogen, resulting in an asymmetric dimethyl-arginine (ADMA/Rme2a)^[6]^. Type II PRMTs also catalyze the synthesis of MMA and can add a second methyl group at the terminal nitrogen adjacent, resulting in symmetric dimethyl-arginine (SDMA/Rme2s)^[6]^. Protein lysine methyltransferases (PKMTs) catalyze the addition of up to three methyl groups at the ε-amino group of a lysine residue (Kme1, Kme2, and Kme3). Most known human PRMTs and PKMTs can methylate histone and non-histone proteins^[7]^ and are potential drug targets due to the fundamental roles in cell biology^[8]^.

The kinetoplastid protozoan parasite *Trypanosoma cruzi* is the etiologic agent of Chagas disease^[9]^, an illness that is estimated to affect about 6 to 7 million people worldwide^[10]^ and which presents a significant burden on the health-care system^[11]^. No vaccine is available, and treatment is carried out with inefficient and highly toxic drugs^[12]^. In addition, *T. cruzi* is a parasite that has a complex life cycle (like other trypanosomes, such as *T. brucei* and *Leishmania* sp.) with different stages of development and different hosts, making it an interesting organism to study the biological impact of PTMs. Recently, the presence of a variety of PTMs in *T. cruzi* histones has been demonstrated, with acetylation and methylation being the most abundant types in these proteins^[13,14]^.

Although large-scale non-histone methylation analysis has been performed on higher eukaryotes^[15-19]^, this is not the case for protozoan parasites, and the function of arginine and lysine methylation in these organisms is still poorly understood. In parasites that evolved from an earliest eukaryotic lineage, such *Giardia duodenalis* (diplomonads), methyl-arginines (and arginine-methyltransferases) are absent^[20]^. In trypanosomatids, *Leishmania donovani*, only 19 methylated proteins were identified, including RNA binding proteins (RBPs), ribosomal proteins, and elongation factor 1-alpha^[21]^. In *L. major*, various RBPs have been shown to be target of PRMT7^[22,23]^, indicating the impact of arginine methylation in the RNA metabolism of this parasite. Cruz et al., investigating the impact of knockout of PRMT7 showed, in murine model, a negative correlation between the levels of *L. major* PRMT7 and parasite pathogenicity^[24]^. Read et al. demonstrated that *T. brucei* PRMT1 knockout, on the other hand, decreases parasite virulence, affect energy metabolism and the starvation stress response^[25]^.

In *T. brucei*, approximately 10% of the proteome contains methyl-arginines, potentially impacting diverse cellular pathways in this parasite^[26,27]^, and, recently, 92 and 70 methyl-lysine sites were reported in *T. brucei* and *T. evansi*, respectively^[28]^. For other parasites such *Plasmodium falciparum*, arginine^[29]^ and lysine^[30]^ methylated proteins are showed involved in diverse biological pathways, including RNA metabolism, protein synthesis, transport, proteolysis, protein folding, and chromatin organization. Here, we applied a bottom-up proteomic approach and bioinformatics analysis to comprehensively characterize the proteome-wide methylation in *T. cruzi*. We identified more than a thousand methyl sites in proteins involved in many biological functions relevant to parasite, shedding light on the importance of methylation beyond histone proteins in *T. cruzi*, and providing another promising landscape for understanding the biology of this pathogen.

## 2. Materials and Methods

### 2.1 Cell culture

*Trypanosoma cruzi* Dm28c epimastigotes in the exponential growth phase were cultured in Liver Infusion Tryptose (LIT) medium^[31]^ supplemented with 10% fetal bovine serum and incubated without agitation at 28 °C. Epimastigotes at the exponential growth phase were obtained in four biological replicates (R1, R2, R3, and R4) in the order of 1.5 ×10^9^ cells.

### 2.2 Protein extraction, separation, and digestion

Cells were washed in PBS, resuspended in lysis buffer (4% SDS, 100 mM Tris-HCl pH 7.5, 100 mM DTT) (240 μL/3 × 10^8^ cells), vortexed for 30 seconds, heated for 3 min at 95 °C and sonicated for 1 hour at room temperature. To remove debris, samples were then centrifuged at 20,000*g* for 5 min at 20 °C, and the supernatant was transferred to a clean tube. The samples were quantified using Qubit fluorometer according to the manufacturer’s instructions (Invitrogen) and Bradford assay^[32]^. *T. cruzi* protein extract (25-30 ug/replicate) was separated by SDS-PAGE (4 - 20% acrylamide) and stained with Coomassie Blue G-250, and destained without organic solvent^[33]^. Each gel lane was sliced horizontally into 10 fractions covering the different molecular weight ranges, and each fraction was cut into 1 mm^3^ cubes, which were transferred to a clean microfuge tube and submitted to in-gel digestion as described elsewhere^[34]^. Briefly, the gel pieces were destained twice with 25 mM ammonium bicarbonate (ABC) in 50% ethanol, dehydrated with 100% ethanol, reduced with 10 mM DTT in 50 mM ABC, alkylated with 55 mM iodoacetamide in 50 mM ABC, and digested with 12.5 ng/µL trypsin in 50 mM ABC and incubated for 16h at 37°C. Digestion was stopped by adding trifluoroacetic acid (TFA) to a final concentration of 0.5%. Peptides were extracted twice with 30% acetonitrile (MeCN) in 3% TFA and twice with 100% MeCN, then dried in a Speed Vac and desalted using C18 StageTips^[35]^ prior to nanoLC-ESI-MS/MS.

### 2.3 NanoLC-ESI-MS/MS analysis

Tryptic peptides of four biological replicates were separated by online reverse phase (RP) nanoscale capillary liquid chromatography (nanoLC) and analyzed by electrospray mass spectrometry in tandem (ESI-MS/MS). The samples were analyzed at the mass spectrometry facility RPT02H at Carlos Chagas Institute/Fiocruz-Parana with an LTQ-Orbitrap XL ETD or an Orbitrap Fusion Lumos mass spectrometer (Thermo Fisher Scientific). For LTQ-Orbitrap, peptide mixtures were injected in an Easy-nLC 1000 system (Thermo Fisher) with a 30 cm analytical column (75 μm inner diameter) in-house packed with C18 resin (ReproSil-Pur C18-AQ 2.4 μm), eluted from 5 to 40% MeCN in 5% DMSO and 0.1% formic acid in a 120 min gradient. The nanoLC column was heated at 60 °C to improve chromatographic resolution and reproducibility. Peptides were ionized by nanoelectrospray (2.7 kV) and injected into the mass spectrometer. Mass spectra were obtained by Data-Dependent Acquisition (DDA), with an initial Orbitrap scan with resolution R = 15,000 followed by MS/MS of the 10 most intense ions in LTQ (Iontrap). These precursor ions were fragmented by Collision-Induced Dissociation (CID) with normalized collision energy = 35%, activation time of 30 ms and activation q = 0.25. Singly charged precursor ions were rejected. Parallel to the MS2, a full scan was conducted in Orbitrap with a resolution of 60,000 (mass range m/z 300-2000), and selected ions were dynamically excluded for 90 seconds. The lock mass option^[36]^, in the presence of DMSO peaks^[37]^, was used in all full scans to improve the mass accuracy of precursor ions. For Orbitrap-Fusion Lumos, peptides were injected in an Ultimate 3000 RSLCnano liquid chromatograph (Thermo Scientific) with an analytical column of 15 cm with 75 um of internal diameter, containing C18 particles of 3 micrometers in diameter. The gradient elution was 0.1% formic acid (phase A) and 0.1% formic acid, 95% acetonitrile (Phase B). Flow rate 250 nL/min. Linear gradient from 5 to 40% acetonitrile in 60 min. In the ion source, the peptides were ionized by nanoelectrospray (2.3 kV) with ion transfer tube temperature of 175 °C. The MS1 Orbitrap resolution of 120,000. Included precursor ions with 2-7 charge state. Maximum injection time of 50 ms. Selected ions were dynamically excluded for 60 seconds. In the MS2, the precursor ions were fragmented by Higher energy Collisional Dissociation (HCD) with collision energy = 30%, maximum injection time of 22 ms, with an Orbitrap resolution of 15,000.

### 2.4 Search and identification of protein

Raw data of four biological replicates from the LC-MS/MS analysis was analyzed in the MaxQuant platform^[38]^, version 2.2.0.0. The proteins were identified using a database containing a total of 19,242 protein sequences of *T. cruzi* CL Brener, downloaded on november 22, 2022, from UniProt (www.uniprot.org). Contaminants (human keratins, BSA, and porcine trypsin) were added to the database, as were their reverse sequences. The search for methylated sites used the following criteria: MS tolerance of 20 ppm (Orbitrap), MS/MS tolerance of 0.5 Da (Iontrap), allowing for two missed cleavages, the peptides length searched for with at least seven amino acids, and trypsin with specific cleavage. Carbamidomethylation of cysteines was determined as a fixed modification. Monomethylation and dimethylation of lysines/arginines, and trimethylation of lysines were searched as variable modifications, as were methionine oxidation and N-terminal acetylation. The match between runs algorithm was used. A false discovery rate (FDR) of 1% was applied to the protein, peptide, and methyl site levels.

### 2.5 Bioinformatics analyses of *T. cruzi* methylproteome

The functional classification and enrichment analyses of Gene Ontology (GO)^[39]^, and Kyoto Encyclopedia of Genes and Genomes (KEGG)^[40]^ were performed using the Database for Annotation, Visualization, and Integrated Discovery (DAVID) tool^[41]^ version 2021 (Dec. 2021) with count 2 and EASE 0.1 and also through TriTrypDB^[42]^ using P-value cuttof 0.05. Protein consensus sequence analyses were performed using *k*pLogo^[43]^ and iceLogo^[44]^ with the *T. cruzi* proteome as a reference set. In the Venn analysis was used jvenn tool^[45]^. The protein-protein interactions were visualized using the Search Tool for the Retrieval of Interacting Genes/Proteins (STRING)^[46]^ in Cytoscape (version 3.9.1)^[47]^ with confidence score cutoff of 0.70 for protein interactions.

## 3. Results

### 3.1 GeLC-MS/MS analysis reveals abundant arginine and lysine methylation in *T. cruzi* epimastigotes

We applied a gel-based proteomic approach to characterize the methylproteome of arginine and lysine residues of *T. cruzi* epimastigotes with four biological replicates (Fig. 1A). Our analysis of the total *T. cruzi* protein extract resulted in the acquisition of 1,381,247 MS/MS spectra, 309,008 of which were identified when compared to the database. After exclusion of the identified contaminants and reverse sequences, a total of 34,865 peptides and 5,007 proteins were identified. Among them, 824 methylated proteins (Table S1) and a total of 1252 methyl sites (Table S2) were detected, 88.4% with a high-confidence methylation site (localization probability >0.80). The number of total and methylated proteins (Fig. 1B and Table S1) were similar in each replicate, from 15.5% to 16.1% of total proteins was identified methylated. We found that 86.4% (n=712) of total methylated proteins were identified in ≥2 biological replicates (Fig. 1C). Considering ≥3 biological replicates the overlap is 79.5% (n=655) and considering all replicates the overlap of methylproteins is 73.3% (n=604). Results consistent with the reliable identification of sites and peptides by our MS analysis. Similar numbers of methylated sites were identified in each replicate (Fig. 1D and Table S2) considering the sum of K and R methylated sites in each replicate R1 (n=615), R2 (n=610), R3 (n=581), and R4 (514). The R4 had a slightly lower number of identified sites compared to the other replicas, as well as total and methylated proteins. The representative spectra of methylated sites are shown in Fig. S1. Then we analyzed the distribution of these sites per protein, and we found that 44.3% identified proteins have more than one arginine methylation site, and only 1.09% containing >10 methylation sites (Fig. 1E).

**Figure 1.**
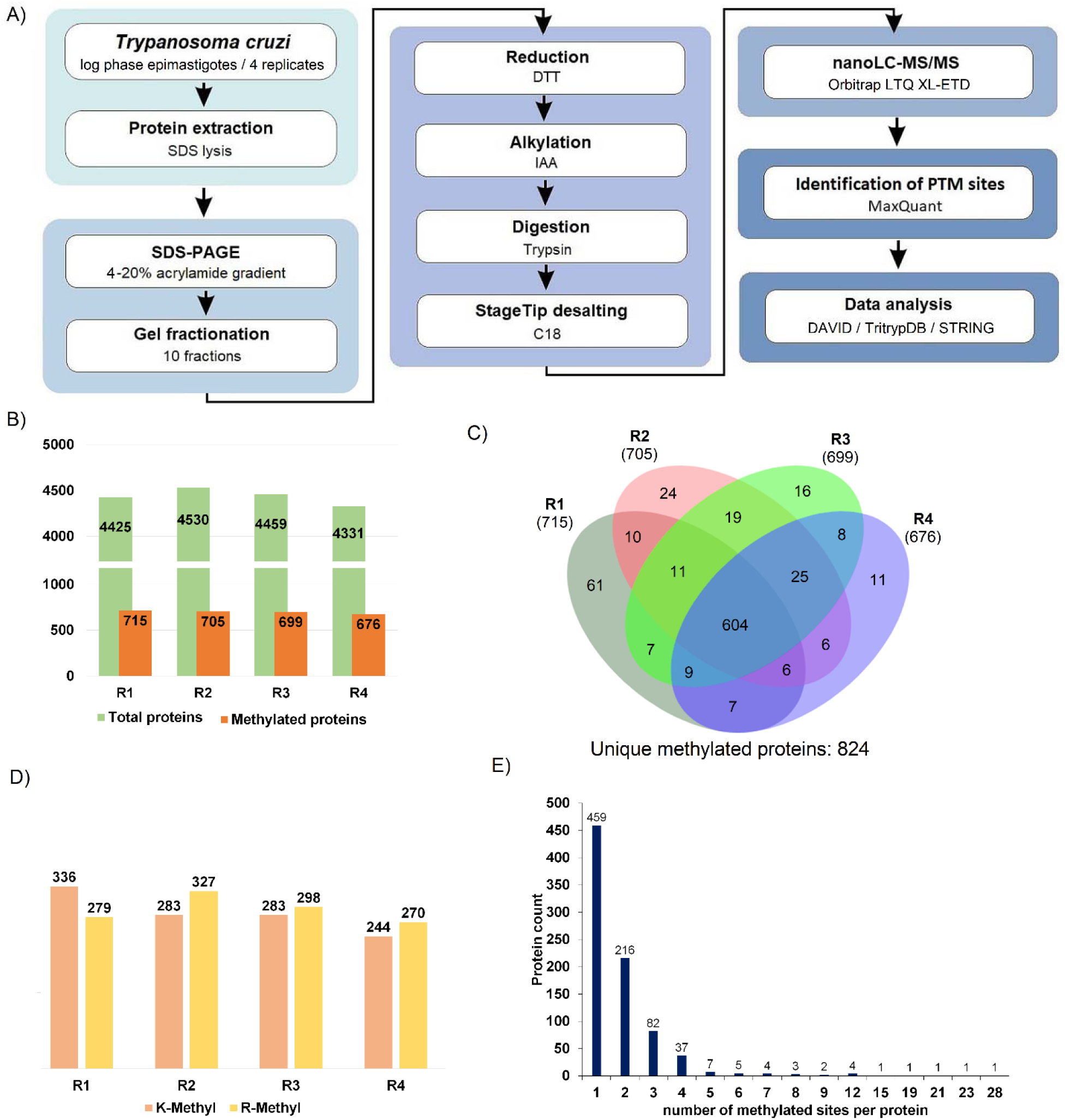
Workflow and global numbers of *T. cruzi* methylproteome. A) Proteomic workflow applied to characterize *T. cruzi* methylproteome. B) Comparative numbers of total and methylated proteins in different biological replicates. C) Venn diagram showing the number of methylated proteins identified different biological replicates. D) Number of methylation sites (K and R) identified in different biological replicates and (E) distribution of arginine methylation sites per protein, considering all methylation types (me, me2 and me3).

### 3.2 Methylated proteins impact numerous processes in *T. cruzi*

In order to identify the processes and pathways potentially regulated by methylated proteins in *T. cruzi* epimastigotes, we performed enrichment analysis of proteins based on Gene Ontology (GO)^[39]^ and KEGG pathways^[40]^ terms available in DAVID^[41]^ and TritrypDB^[42]^ with methylated proteins identified in ≥2 biological replicates. GO analysis revealed a widespread cellular processes and localization of methylated proteins both on nucleus and cytoplasm (Fig. 2 and Table S3). The most enriched biological process terms associated with methylated proteins harboring protein folding (P-value: 2.16 × 10^−7)^ and translation (P-value: 3.65 × 10^−7^) due a high number of ribosome subunits, chaperones and chaperonins over-represented in our results in relation of its GO terms in *T. cruzi* proteome. Methylated proteins were algo associated with different amino acid and carbohydrate metabolisms. These different biological and molecular processes are confirmed when we analyzed the KEGG pathways. In this analysis the glycolysis/gluconeogenesis (P-value: 2.27 × 10^−4^) and ribosomes (P-value: 2.35 × 10^−4^) appears more enriched, followed by biochemistry pathways involved in Krebs cycle and energy generation.

**Figure 2.**
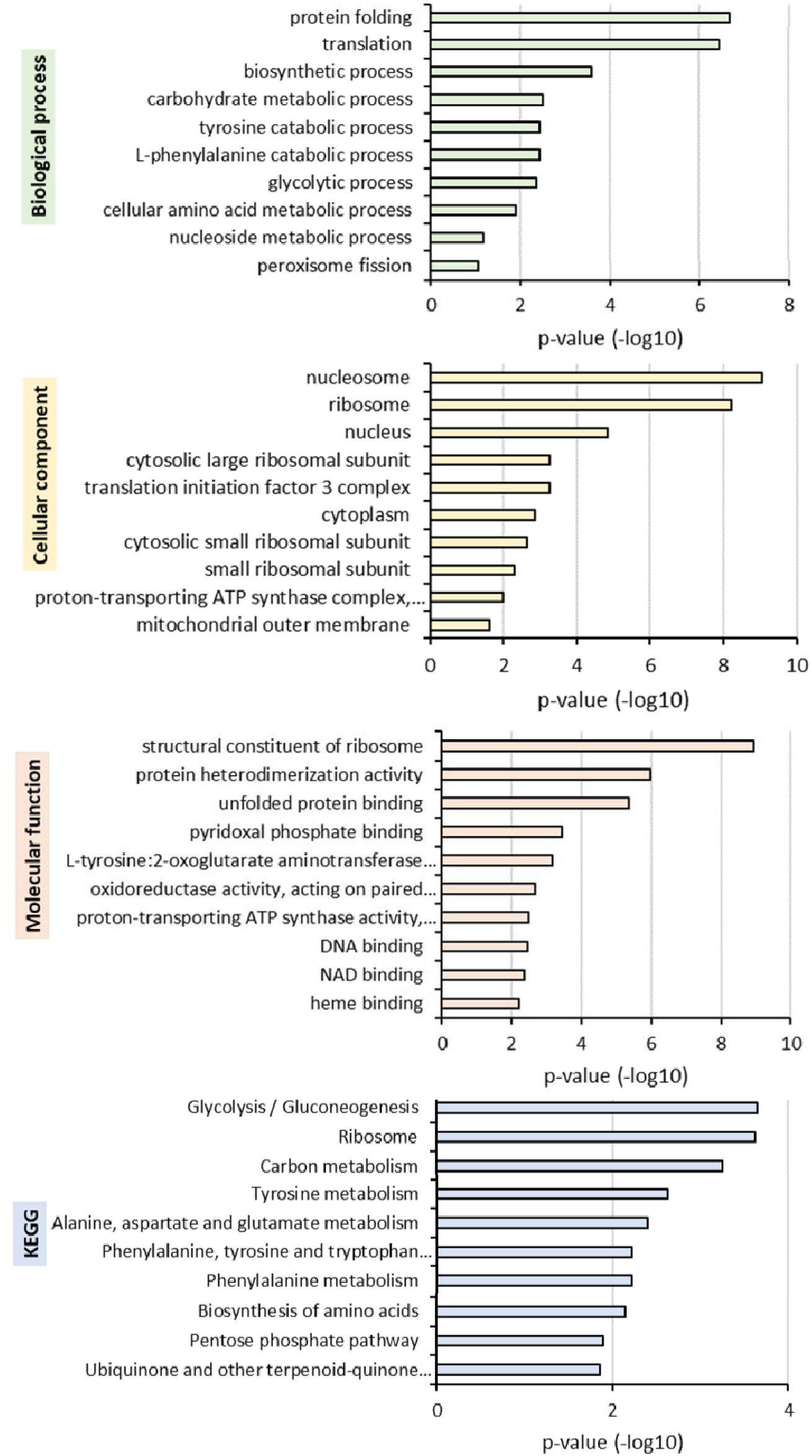
Functional characterization of *T. cruzi* methylproteome. GO-based enrichment analysis of methylated proteins (K and R methylated proteins combined) identified in ≥2 biological replicates (712 methylproteins) according to biological processes, molecular function, cellular component, and KEGG pathways. Top 10 terms more enriched (P-value) in DAVID^[41]^ chart analysis are showed. Detailed results are available in Table S3.

To improve the gene ontology enrichment analysis obtained with the DAVID tool, the datasets were also analyzed in TritrypDB, the database specifically of kinetoplastids parasites, and the results shown a similar enriched terms in each category of GO, with more specifically terms for molecular functions such elongation factors and RNA binding confirming the results associated to translation in DAVID bioinformatic tool (Fig. S2 and Table S3). Furthermore, elongation factors and tubulin proteins were identified with multiple (>10) methylation sites (Table S2).

### 3.3 Lysine- and arginine-methylated proteins show different enriched biological themes

Intrigued with the occurrence of methylation in similar numbers both in lysine (K) and arginine (R) (Fig. 1D), we evaluate whether different cellular functions are regulated by K-Me and R-Me proteins in *T. cruzi*. We performed functional enrichment analysis with the subproteomes separately using DAVID functional enrichment tool considering proteins identified in ≥2 biological replicates (Fig. S3). Many GO terms appear enriched in both subproteomes, although with different enrichment values. Comparing biological processes, the most enriched theme was *protein folding* for both subproteomes, more enriched in K-Me (P-value: 3.92 × 10^−5^) than in R-Me (P-value: 1.99 × 10^−4^) proteins. Interestingly, the *translation* process appears more enriched in R-me (P-value: 8.68 × 10^−4^) than in K-me (2.72 × 10^−2^), with more than two orders of magnitude. Comparing the localization of methylated proteins, some terms are exclusively of each subproteome, such *nucleosome* appears highly enriched in R-Me proteins (P-value: 1.86 × 10^−9^), while *mitochondrial outer membrane* (P-value: 7.31 × 10^−3^) in K-Me. Molecular function related to GTPase activity (P-value: 2.79 × 10^−5^) and GTP binding (P-value: 5.26 × 10^−4^) are more enriched in K-Me proteins, harboring GTP-binding and different elongation factors proteins. in the enrichment of KEGG pathways, only R-Me presents *ribosome* enriched (P-value: 8.32 × 10^−3^), according to GO categories.

### 3.4 Protein-protein interaction of functional clusters provides a network view of the methylproteome

In order to evaluate the protein-protein interaction (PPIs) of methylated proteins functional related in *T. cruzi*, we searched for functional enriched clusters in DAVID annotation clustering tool and mapped the interactions of proteins available in STRING database. Considering methylated proteins identified in ≥2 replicates (n=712), the DAVID analysis resulted 11 enriched clusters and a subset of 169 proteins present interaction annotation in the STRING database. The significantly enriched protein clusters included translation/ribosome, translation initiation factors, nucleosome and DNA binding, ATP/GTPase activity and elongation factors, glycolysis/gluconeogenesis and carbon metabolism, amino acid and carbohydrate metabolisms (Fig. 3 and Table S5). Proteins bearing K or R methylation (or both) interact with each other, and no protein interaction cluster bearing only one type of methylation were observed. The clusters with more interaction partners is related to translation and glycolysis/gluconeogenesis and carbon metabolism, important pathways in *T. cruzi*. For example, among translational initiation factors cluster, we identified the eukaryotic translation initiation factor 3 subunit 7-like protein (Q4DHE0) and subunit 8 (Q4DQJ3), two putatives translational factors (Q4DSL1 and Q4E030), and two “uncharacterized proteins” (Q4DDK1, and Q4DL69). The nucleosome and DNA-binding cluster contains histones H2A (Q4DYB1), H2B (Q4CTD8), H3.V (Q4DRD1), DNA topoisomerase IA (Q4DE64) and IB (Q4DTW1), DNA helicase (Q4E255), DNA ligase (Q4DX91), and DNA-directed RNA polymerase subunit (Q4DDX5). Collectively, these interaction data demonstrate that methylation is present in important protein complexes in the parasite.

**Figure 3.**
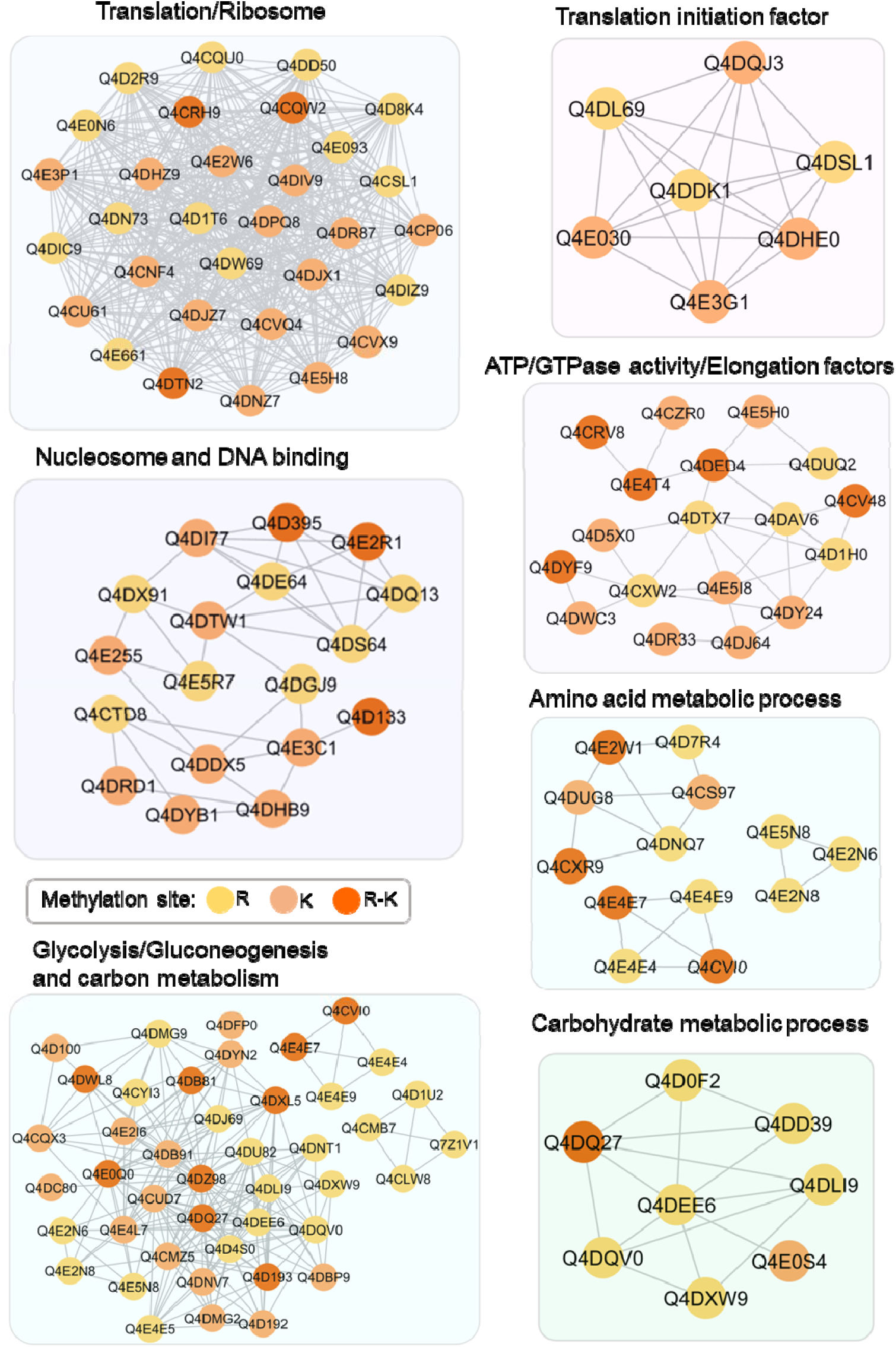
Interaction clusters of methylated proteins in *T. cruzi*. The gene ontology and KEGG pathways of methylated proteins were searched in DAVID, and significantly enriched protein clusters were visualized using STRING^[46]^ and Cytoscape^[47]^. The proteins with high-confidence interactions (score ≥0.70) were included. Data are representative of ≥2 biological replicates. Detailed data are listed in Table S5.

### 3.5 *T. cruzi* arginine and lysine methyl sites are surrounded by a different amino acid pattern

In order to identify amino acid patterns surrounding methyl sites in *T. cruzi* and find possible census motifs of PRMTs and PKMTs in parasite we investigated the frequencies of the amino acid surrounding sites. Despite the similar distribution of total lysine (K) and arginine (R) methylated sites of *T. cruzi* methylproteome (625 and 627, respectively) different amino acids pattern occur more frequently in the vicinity of arginine and lysine (Fig. 4). The set of all methyl-lysine sites are enriched upstream and downstream the site for glutamic acid (E), due the presence of more sequences of Kme2 with these patterns. Interesting, in the lysine monomethylation (Kme) sequences the most significant pattern upstream is three cysteines (C) the same pattern found in arginine monomethylation (Rme), and these, with the highest number of sequences and therefore even more enriched. The most significant motifs (k-mer) at each position are detailed in Table S6.

**Figure 4.**
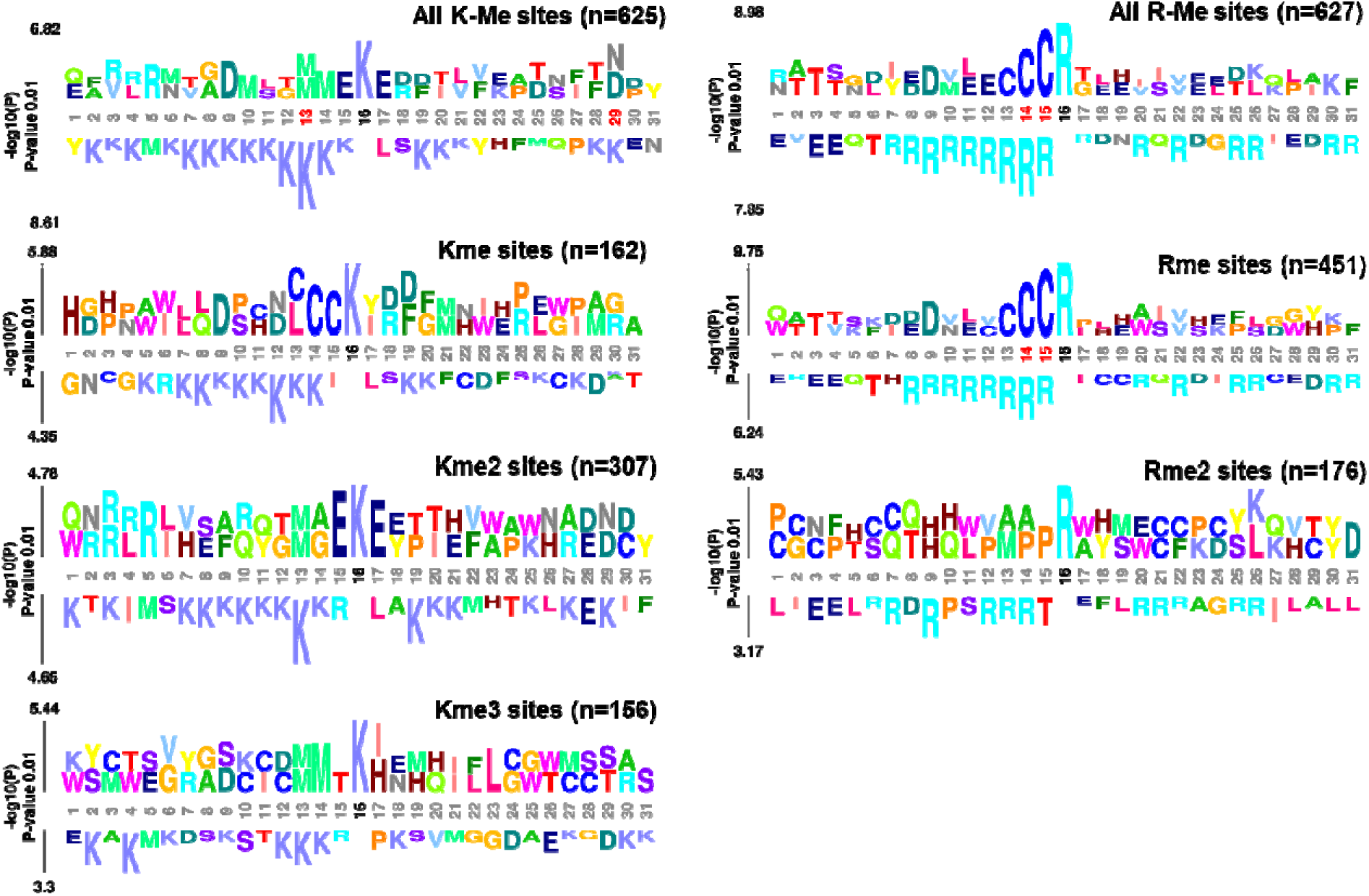
Over-represented amino acids surrounding *T. cruzi* methylated sites. Sequence window of all K-Me and R-Me sites and subsets of methylation types (me, me2, and me3) were analyzed using kpLogo^[43]^. K-Me and R-Me present different amino acids patterns surrounding site. The sequence windows with 15 amino acids upstream and downstream of the methylated site were aligned and the probability of the occurrence of amino acids in each position calculated and the most significant is plotted vertically with the total height scaled to its P-value (log10 transformed). Stacked residues at a position represent the most significant (P-value cutoff 0.01) starting (or ending) at this position. The total height is scaled relative to the significance of this motif. At each position, two motifs are shown, the most enriched on the top, and the most depleted on the bottom. Positions containing residues with Bonferroni corrected P-value smaller than 0.01 are highlighted by red. The most significant motifs (k-mer) at each position are detailed in Table S6.

### 3.6 *T. cruzi* methylated proteins presents co-occurrence with phosphorylation

The interplay of methylation and phosphorylation has already been reported in human^[48,49]^, however this has not yet been verified in *T. cruzi*. In view of this, we compare our methylproteome results with previous large-scale studies describing phosphorylation in *T. cruzi*^[50-52]^, and through a direct comparison, we found that of the 712 methylated proteins, we found that 171 (24%) reportedly contain phosphorylated sites. The GO enrichment analysis in TritrypDB reveals 15 enriched terms, mostly related to parasite flagellum and cytoskeleton, GTPase activity, ion binding, and RNA-binding. The most enriched (P-value: 5.94 × 10^−9^) was proteins localized in flagellum and cytoskeleton of parasite, such paraflagellar rod protein, intraflagellar transport protein, and flagellar attachment zone protein. The phosphorylation and dephosphorylation of paraflagellar rod protein in *T. cruzi* seems to be involved with the adhesion of trypomastigote form previously to cell infection^[53]^. Trypanosomes have a relevant capacity of modulates mRNA abundance and translational repression by RNA-binding proteins (RBPs) and the RBPs themselves are regulated by PTMs, including phosphorylation and methylation^[54]^. We found proteins related to RNA-binding enriched in multiples terms (translation factor activity, P-value: 1.03 × 10^−2^; RNA binding, P-value: 1.91 × 10^−2^ and mRNA metabolic process, P-value: 3.49 × 10^−2^) includes RNA-binding proteins, eukaryotic translation initiation factor 3 and 4A-1, elongation factors 2 and TU (mitochondrial), RNA capping enzyme, and U2 splicing auxiliary factor. Proteins related to GTPase activity, ion binding, peptidase activity, carbohydrate metabolic process, and amino acid metabolic process were also well represented.

## 4. Discussion

Post-translational Modifications (PTMs) have a regulatory role in many essential cellular processes that range from gene transcription to signal transduction, including protein methylation^[4]^. Methylation in non-histone proteins plays a role in regulating cell signaling, impacting pathophysiological processes, including cancer^[7,55-57]^. In trypanosomes, few large-scale studies have been conducted to elucidate the proteome-wide impact of PTMs by non-histone proteins. Recently, Zhang et al.^[28]^ performed a broad search of PTMs in African trypanosomes, *T. brucei* and *T. evansi* showing that at the level of PTMs these two phylogenetically close parasites have important differences. Differences were also observed between *T. brucei* and *T. cruzi* when their acetylomes were compared^[58]^. Another PTM studied in trypanosomes is phosphorylation, investigated in *T. brucei*^[59,60]^ it was identified modulating RNA binding proteins in co-ordinating with heat shock proteins^[59]^, and in *T. cruzi*^*[50-52]*^, phosphorylation is modulated in metacyclogenesis, impacting translation, oxidative stress, and the metabolism of protein, lipids, and carbohydrates^[50]^. Protein-wide methylation study in trypanosomes was conducted by Read et. al.^[27]^, showing that arginine methylation impact different processes in *T. brucei*. However, to date, *T. cruzi* lacks studies on methylation at a proteome-wide level. To fill this gap, we applied a large-scale proteomic workflow to identify and mapping arginine and lysine methylated proteins in *T. cruzi* epimastigotes. Using a well-established GeLC-MS/MS approach^[34]^ with four biological replicates, we identified 1253 methyl sites in 824 methylated proteins, of these, 712 proteins were identified in two or more biological replicates. These methylated proteins are in different cell compartments and present a wide range of functions, demonstrating the influence of methylation on several biological processes of the parasite, including translation, protein folding, amino acid and carbohydrate metabolism, RNA metabolism, among others important processes. Our protein-protein interactions analysis of functional enriched proteins reveals that important complex with robust interactions data are impacted by methylation and present different levels of methylation in arginine, lysine, or both. Among the methylated proteins detected in *T. cruzi* are the related to translation processes such ribosomal proteins and RBPs. RBPs are known substrates of PRMT in different organisms^[61-63]^ and are fundamental to differential gene regulation processes in *T. cruzi*, due to the post-transcriptional events regulating the gene expression^[64]^. We identified proteins involved in RNA processing with methyl sites, similarly, to demonstrated in *T. brucei* procyclic form impacted by arginine methylation^[27]^. While in *T. brucei*, the main represented functions were related to the cytoskeleton and locomotion, here, on the other hand, the main function of the majority of identified methylated proteins identified was translation and protein folding. Indeed, the same PTM can affect different cell processes; this dissimilarity can be explained by the differences in parasite’s biology at a PTM level. For example, recently, Schenkman et al.^[58]^, analyzing lysine acetylation in trypanosomes, revealed that protein acetylation is involved in a very distinct set of acetylated proteins when comparing *T. cruzi* and *T. brucei*. Therefore, these functional differences between methylproteomes (and acetylomes) are compatible to with the broad role of PTMs in cell signaling already demonstrated in other eukaryotes, and that seems to also exist between trypanosomes^[65]^.

Although our work was not directed towards enriched histone fractions, we identified methylated sites in different histones: H4R90me2, H4R54me2, H3.VK95me3, H2BK97me, H1K10me2, H2BR61me, H2BR61me2, and H2AK102me2. The methylated histones H3.VK95me3 (Q4DRD1), H2BK97me (Q4DHB9), H2BR61me2 (Q4CTD8), and H2AK102me2 (Q4DYB1) were mapped n our interaction cluster analysis.

Another important feature of methylation is the target motifs of methyltransferases. Our analysis of amino acid patterns surrounding methylation sites revealed different residues in K-Me and R-Me. The vicinity of all lysine methylated sites are mostly enriched for glutamic acids, also seen in *P. falciparum*^[29,30]^ and *G. duodenalis*^*[20]*^, and arginine methylated sites are enriched upstream for cysteines. Well-defined patterns, such as “RGG,”, are not always the preferential substrate for PRMTs, as demonstrated for human PRMT1, it was suggested that residues upstream to the modification site are also important in substrate recognition^[66]^. As in humans, PRMT1 is the main arginine methyltransferase in *T. brucei*^[67]^ and *Toxoplasma gondii*^*[68]*^. *T. cruzi* PRMT1 have the methyltransferase domain region is well conserved with the homologous PRMT1 in *T. brucei* and human, especially in residues that promote the binding of the enzyme to the molecules of AdoMet. In view of this, our data for *T. cruzi* reinforces that the target motifs of methyltransferases seem to be broader.

Methylated proteins also impact important processes like RNA processing, which is responsible for the assembly of multiprotein complexes on primary transcripts, mature mRNAs, and stable ribonucleoprotein components of the RNA processing machinery^[69]^. Recently, Amorim and colleagues^[50]^ demonstrated that EF-1-α and EF-2 are phosphorylated in *T. cruzi* during the final phase of metacyclogenesis. When we compared the methylproteome data with phosphoproteome data previously reported, methylated proteins involved with RNA processing also have phosphosites. Here, we identified RNA-binding proteins, eukaryotic translation initiation factor 3 and 4A-1, elongation factors 2 and TU (mitochondrial), RNA capping enzyme, and U2 splicing auxiliary factor. Furthermore, elongation factors and tubulin were identified with multiple (>10) methylation sites. These elongation factors are key pieces of the translation process and, may act beyond the canonical process in eukaryotes, including in *T. cruzi*^[70]^. They may therefore be used as potential drug targets, as is the case for EF-2 in *P. falciparum*^[71]^.

An important class of mediators of signal transduction are kinase proteins. We detected methyl sites in 33 proteins annotated with kinase function, among them serine/threonine protein kinase, phosphatidylinositol 4-phosphate 5-kinase, adenosine kinase, arginine kinase, phosphoglycerate kinase, pyridoxal kinase, glycerate kinase, pyruvate dikinase, among others. Interestingly, serine/threonine-protein kinase (Q4CXF6) present gene ontology of transmembrane and integral membrane component. Kinases, if located on the surface of trypanosomatids, can phosphorylate host molecules or parasites to modify their environment^[72]^.

The interplay of different PTMs is likely to regulate multiple cellular processes^[65]^, both in histone^[73]^ and non-histone proteins^[74]^. It is known, for example, that arginine methylation affects other methylated residues, such as lysines themselves^[75]^, and other modifications, such as acetylation and phosphorylation^[73]^. In addition, there is an interrelationship between the methylation of lysines 4 and 79 of histone H3 (H3K4 and H3K79) with the ubiquitination of lysine 123 of histone H2B (H2BK123) in *Saccharomyces cerevisiae*, where the non-ubiquitination of K123 prevents methylation of K4 and K79^[76]^. In mammals, an antagonism between arginine methylation and serine phosphorylation in the C-terminal domain of RNA polymerase II impacts the transcription of specific genes^[77]^ and methylation prevents recognition by a cyclin-dependent kinase (Cdk) in serine residue, preventing the phosphorylation of this substrate, controlling the progression of the cell cycle^[48]^. We found that 171 methylated proteins were previously reported with phosphorylation sites^[50-52]^. This leads to the hypothesis that there may be a potential crosstalk between these modifications working to adjust the function of the protein in a tertiary structure level that influence the interaction with other binding molecules, or they may be further away to prevent interference, so that modifying enzymes can independently bind to the respective sites^[78]^.

Our methylproteome results with the analysis of previously reported phosphoproteome data provide candidates to study of potential crosstalk between methylation and phosphorylation in *T. cruzi*. Further characterization is necessary to identify a methyl-phosphorylation switch coregulation in this parasite with important role in cell cycle and DNA damage repair, as recently demonstrated in mammals^[79]^. This is necessary to understanding the signaling of PTMs at a proteome level, even more in organisms with unique features such as trypanosomes.

Collectively, our data gave another status for protein methylation in the biology of *T. cruzi* and indicated that it has great potential to have important cell regulation roles, and could act in interplay with another PTMs, such phosphorylation. Our data show that protein methylation is present in many processes in *T. cruzi* and that this modification should be further investigated, to reveal key parts in the biology of this parasite and potential chemotherapeutic candidates.

## 5. Associated Data

The mass spectrometry proteomics data have been deposited to the ProteomeXchange Consortium via the PRIDE^[80]^ partner repository with the dataset identifier PXD034140.

## Acknowledgments

The authors thank Fiocruz for using the Technological Platforms Network. Also thank CNPQ, CAPES and Fiocruz for financial support.

## Abbreviations

ABC: ammonium bicarbonate
ADMA: asymmetric dimethyl-arginine
ATP: adenosine triphosphate
BP: biological process
CC: cellular component
COG: Clusters of Orthologous Groups
DTT: 1,4-dithiothreitol
ESI: electrospray ionization
GO: Gene Ontology
HPLC: high-performance liquid chromatography
IAA: iodoacetamide
KEGG: Kyoto Encyclopedia of Genes and Genomes
LC-MS/MS: liquid chromatography with tandem mass spectrometry
MF: molecular function
MMA: monomethyl-arginine
MS: mass spectrometry
MeCN: acetonitrile
PBS: phosphate-buffered saline
PKMT: protein lysine methyltransferase
PRMT: protein arginine methyltransferase
PTM: post-translational modification
Rme1: monomethyl-arginine
Rme2a: asymmetric dimethyl-arginine
Rme2s: symmetric dimethyl-arginine
RBP: RNA binding protein
SAM: S-adenosylmethionine
SDMA: symmetric dimethyl-arginine
SDS: sodium dodecyl sulfate

---

**Table 1.** Functional analysis of proteins with co-occurrence of methylation and phosphorylation in *T. cruzi*

| Category | GO ID | Name | Bgd count | Result count | % of bgd | Fold enrichment | P-value |
| --- | --- | --- | --- | --- | --- | --- | --- |
| CC | 5929 | cilium | 157 | 15 | 9.6 | 6.64 | 5.94E-09 |
| CC | 43226 | organelle | 1520 | 36 | 2.4 | 1.65 | 1.13E-03 |
| MF | 3924 | GTPase activity | 114 | 7 | 6.1 | 4.27 | 1.24E-03 |
| MF | 43167 | ion binding | 2086 | 44 | 2.1 | 1.47 | 2.98E-03 |
| MF | 8233 | peptidase activity | 386 | 12 | 3.1 | 2.16 | 9.47E-03 |
| MF | 8135 | translation factor activity, RNA binding | 128 | 6 | 4.7 | 3.26 | 1.03E-02 |
| BP | 6790 | sulfur compound metabolic process | 70 | 4 | 5.7 | 3.97 | 1.81E-02 |
| MF | 3723 | RNA binding | 425 | 12 | 2.8 | 1.96 | 1.91E-02 |
| BP | 5975 | carbohydrate metabolic process | 110 | 5 | 4.5 | 3.16 | 2.10E-02 |
| BP | 6520 | cellular amino acid metabolic process | 239 | 8 | 3.3 | 2.33 | 2.18E-02 |
| MF | 5198 | structural molecule activity | 254 | 8 | 3.1 | 2.19 | 2.99E-02 |
| BP | 16071 | mRNA metabolic process | 50 | 3 | 6 | 4.17 | 3.49E-02 |
| BP | 6575 | cellular modified amino acid metabolic process | 25 | 2 | 8 | 5.56 | 4.97E-02 |

BP: Biological Process; CC: cellular component; MF: Molecular Function. P-value of 0.05. Bgd: background. Fold enrichment refers to ratio of the proportion of methylproteome genes versus *T. cruzi* genome. P-value (or called EASE score) refers to the significance of gene-term enrichment with a modified Fisher's exact test (EASE score)<sup>[41]</sup>. For detailed results see Table S7.

**Figure S1.**
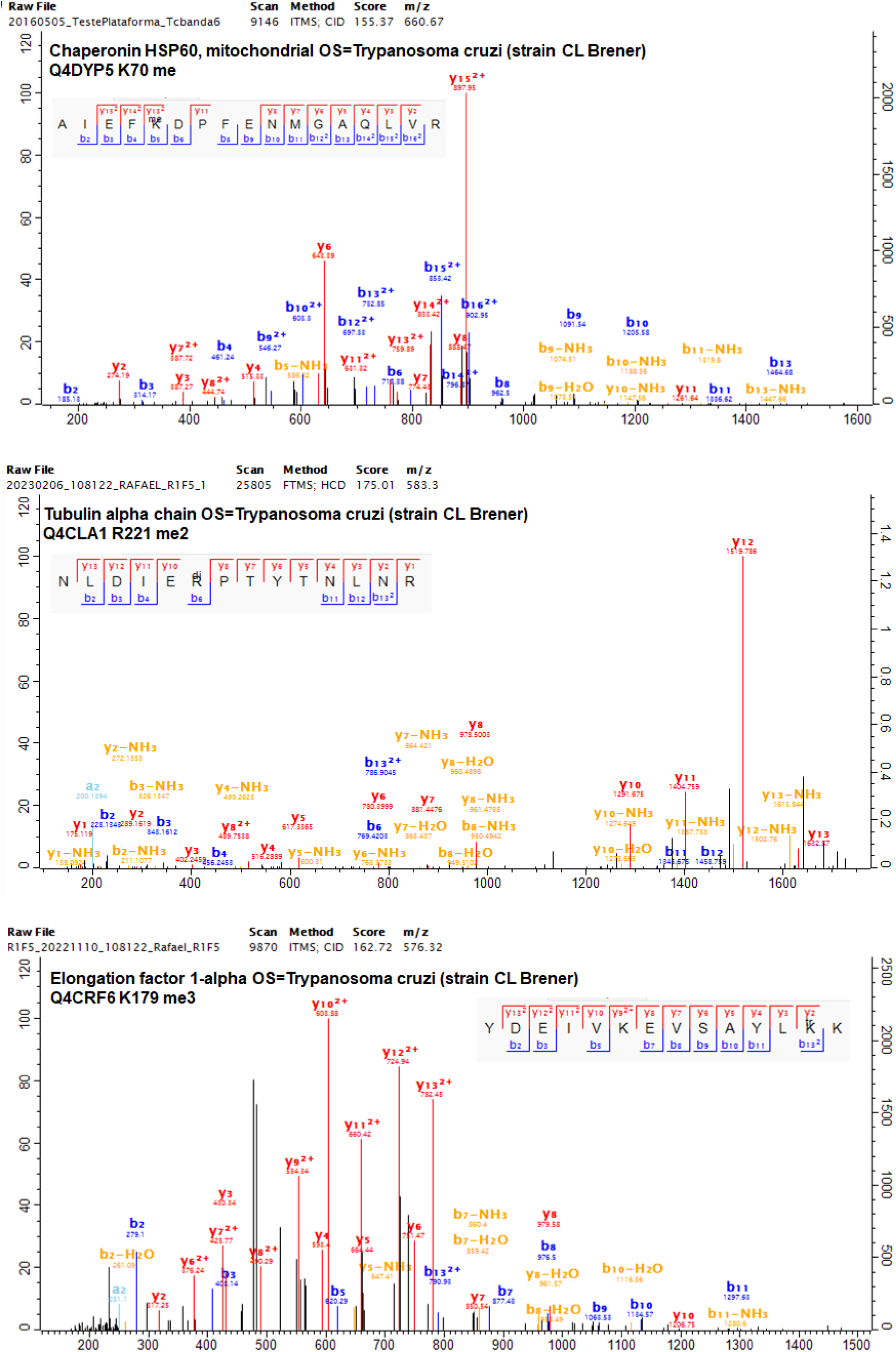
Representative MS/MS spectra of methylated mono-, di-, and trimethylated sites identified in *T. cruzi*. The series of b and y ions are shown flanking the modified site for chaperonin proteins HSP60 (Q4DYP5), tubulin alpha chain (Q4CLA1), and elongation factor 1-alpha (Q4CRF6).

**Figure S2.**
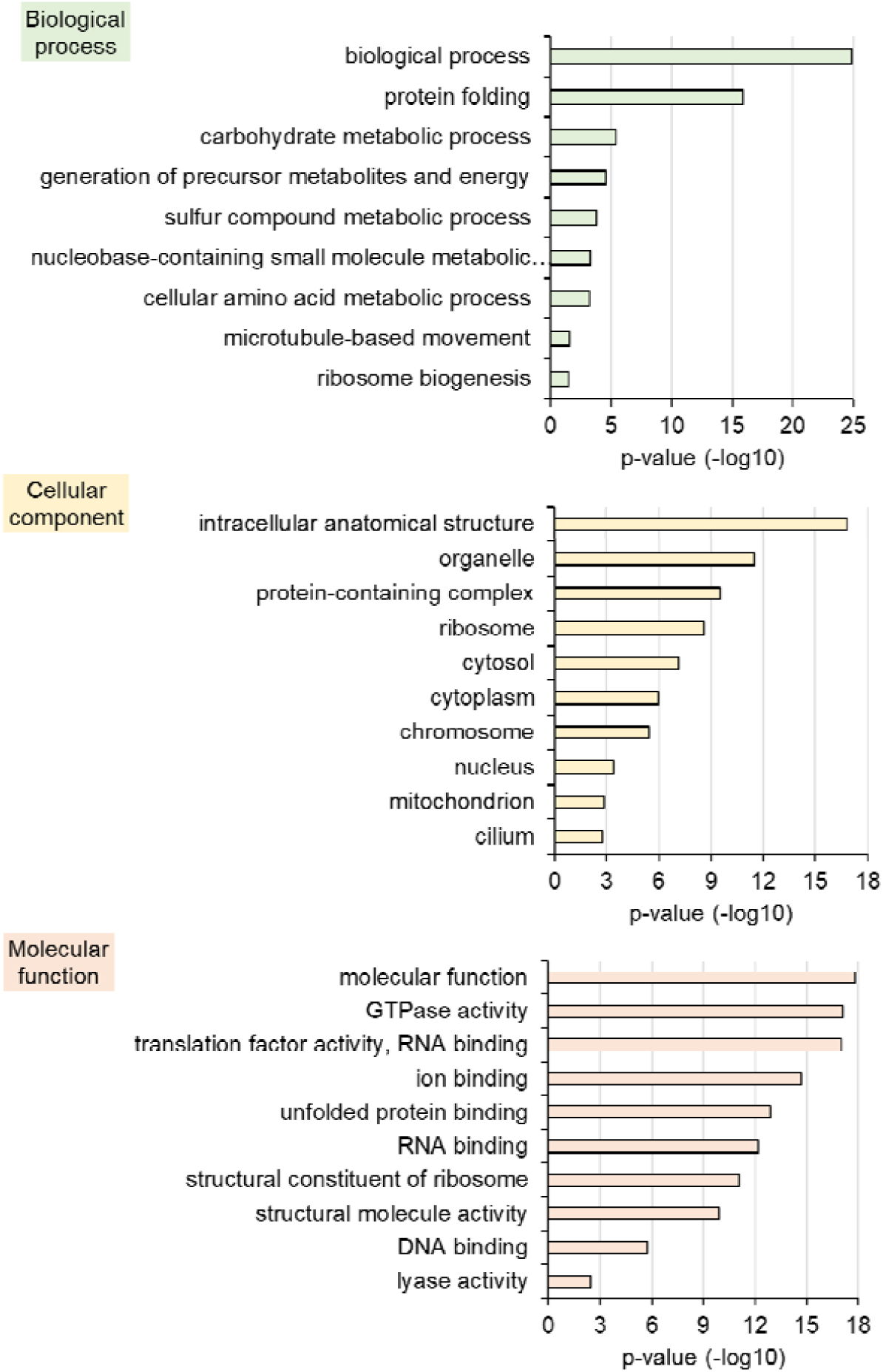
Gene ontology terms enriched in *T. cruzi* methylproteome according to TriTrypDB using a P-value of 0.05 and GO slim terms. For detailed results see Table S3.

**Figure S3.**
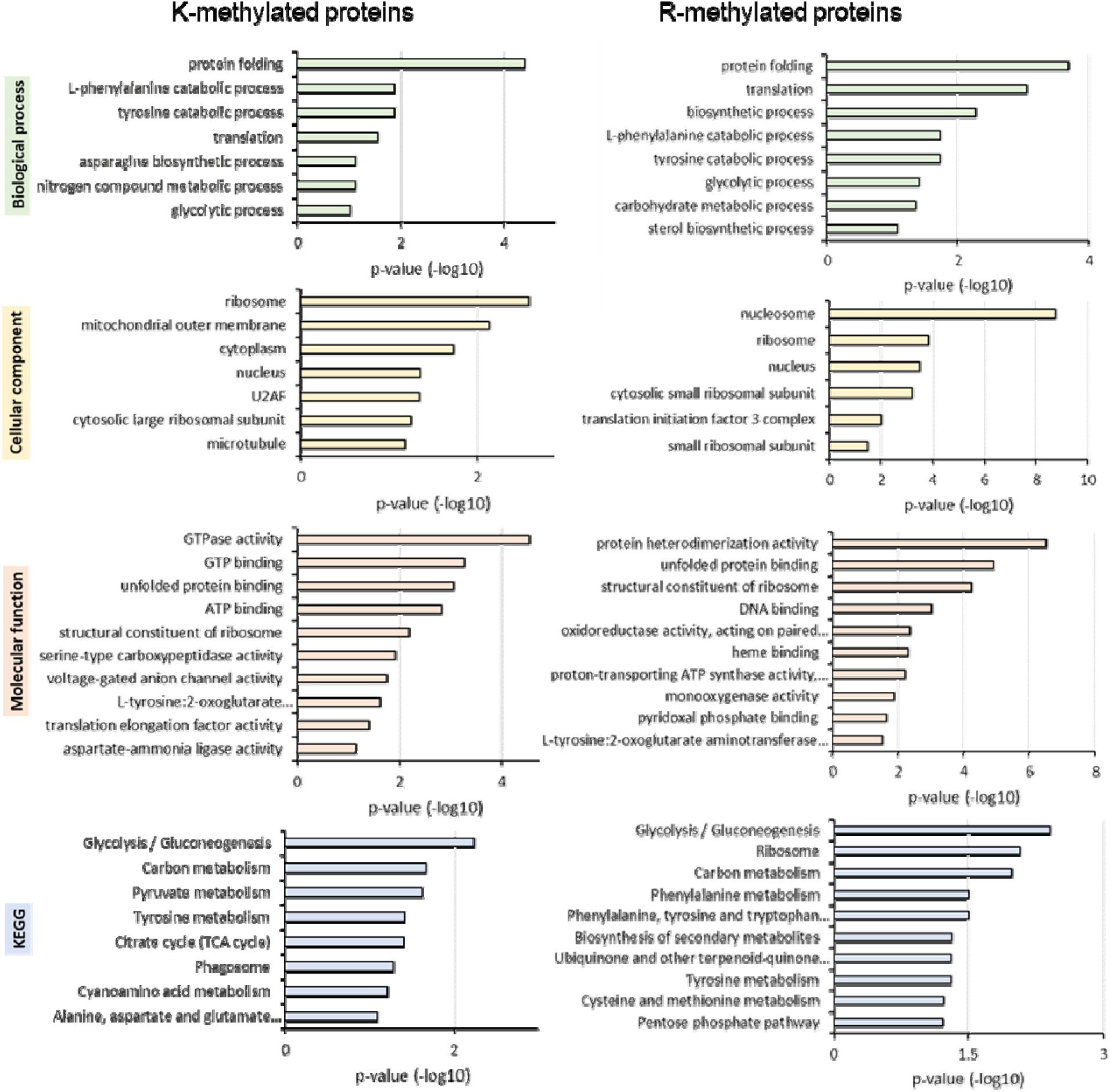
Comparison of functional characterization lysine and arginine methylated proteins. GO-based enrichment analysis comparing proteins methylated in lysine (K) and arginine (R). Were considered proteins identified in ≥2 biological replicates. Top 10 terms more enriched (P-value) in DAVID chart analysis are showed according to biological processes, molecular function, cellular component, and KEGG pathways.

